# Two distinct stimulus-locked EEG signatures reliably encode domain-general confidence during decision formation

**DOI:** 10.1101/2023.04.21.537831

**Authors:** Martina Kopčanová, Robin A. A. Ince, Christopher S. Y. Benwell

**Affiliations:** Division of Psychology, School of Humanities, Social Sciences, and Law, University of Dundee, Dundee, DD1 4HN, UK; School of Psychology and Neuroscience, University of Glasgow, Glasgow, G12 8QB, UK

## Abstract

Decision confidence, an internal estimate of how accurate our choices are, is essential for metacognitive self-evaluation and guides behaviour. However, it can be suboptimal and hence understanding the underlying neurocomputational mechanisms is crucial. To do so, it is essential to establish the extent to which both behavioural and neurophysiological measures of metacognition are reliable over time and shared across cognitive domains. The evidence regarding domain-generality of metacognition has been mixed, while the test-retest reliability of the most widely used metacognitive measures has not been reported. Here, in human participants of both sexes, we examined behavioural and electroencephalographic (EEG) measures of metacognition across two tasks that engage distinct cognitive domains – visual perception and semantic memory. The test-retest reliability of all measures was additionally tested across two experimental sessions. The results revealed a dissociation between metacognitive bias and efficiency, whereby only metacognitive bias showed strong test-retest reliability and domain-generality whilst metacognitive efficiency (measured by M-ratio) was neither reliable nor domain-general. Hence, overall confidence calibration (i.e., metacognitive bias) is a stable trait-like characteristic underpinned by domain-general mechanisms whilst metacognitive efficiency may rely on more domain-specific computations. Additionally, we found two distinct stimulus-locked EEG signatures related to the trial-by-trial fluctuations in confidence ratings during decision formation. A late event-related potential was reliably linked to confidence across cognitive domains, while evoked spectral power predicted confidence most reliably in the semantic knowledge domain. Establishing the reliability and domain-generality of neural predictors of confidence represents an important step in advancing our understanding of the mechanisms underlying self-evaluation.

**Significance Statement:** Understanding the mechanisms underlying metacognition is essential for addressing deficits in self-evaluation. Open questions exist regarding the domain-generality and reliability of both behavioural and neural measures of metacognition. We show that metacognitive bias is reliable across cognitive domains and time, whereas the most adopted measure of metacognitive efficiency is domain-specific and shows poor test-retest reliability. Hence, more reliable measures of metacognition, tailored to specific domains, are needed. We further show that decision confidence is linked to two EEG signatures: late event-related potentials and evoked alpha/beta spectral power. While the former predicts confidence in both perception and semantic knowledge domains, the latter is only reliably linked to knowledge confidence. These findings provide crucial insights into the computations underlying metacognition across domains.

## 1. Introduction

Human decisions are accompanied by a sense of confidence in their accuracy which informs metacognitive self-evaluation. However, studies have shown that confidence does not always appropriately reflect accuracy, leading to distorted metacognition (Shekhar & Rahnev, 2021a, 2021b; Song et al., 2011). Indeed, confidence distortions have been suggested to contribute to both sub-clinical (Benwell et al., 2022b; Rouault, Seow, et al., 2018) and clinical (Hoven et al., 2019) psychopathology. Understanding the underlying neurocomputational mechanisms may help us to understand why metacognition is often sub-optimal and facilitate development of novel interventional targets.

An important open question concerns the extent to which confidence relies on domain-general (versus domain-specific) mechanisms. A domain-general account posits that a shared metacognitive resource is employed to evaluate performance across different cognitive domains (de Gardelle & Mamassian, 2014). Conversely, domain-specific metacognition might rely on computations that are unique to each task (Morales et al., 2018). Behavioural evidence to date has been mixed, with some studies indicating domain-generality (including between sensory modalities (Faivre et al., 2018); perception and memory (Mazancieux et al., 2020; McCurdy et al., 2013; Samaha & Postle, 2017)), but others largely suggesting domain-specificity (Ais et al., 2016; Arbuzova et al., 2022; Baird et al., 2013; Fitzgerald et al., 2017; Kelemen et al., 2000); see Rouault, Mcwilliams, et al., 2018 for review). These discrepancies may partly stem from variability in the measures examined, with distinct latent processes underlying metacognitive judgements (Fleming & Lau, 2014). For instance, metacognitive *sensitivity* indexes the degree to which confidence ratings dissociate correct from incorrect decisions, whereas metacognitive *bias* indexes the overall level of confidence reported (regardless of task accuracy). Metacognitive bias shows stronger correlation across domains than sensitivity (Ais et al., 2016; Benwell et al., 2022b; Mazancieux et al., 2020).

Nevertheless, domain-generality measured at the behavioural level does not rule out the existence of distinct neural mechanisms. Indeed, functional magnetic resonance imaging studies suggest that both domain-general and domain-specific confidence signals for perception and memory coexist in the brain (Baird et al., 2013; McCurdy et al., 2013; Morales et al., 2018; Rouault et al., 2023). Electroencephalography (EEG) allows for measurement of metacognitively relevant neural activity with high temporal resolution. Previous studies have shown that the strength of stimulus-locked EEG responses during decision formation, including the central parietal positivity/P3 component, scale with the reported and/or implicit level of confidence (Azizi et al., 2021; Feuerriegel et al., 2022; Fitzgerald et al., 2022; Gherman & Philiastides, 2015, 2018; Herding et al., 2019; Lim et al., 2020; Rausch et al., 2020; Zakrzewski et al., 2019). Additionally, fluctuations in confidence have also been shown to be dependent on both spontaneous and response-locked levels of EEG activity in the alpha-band (8-12Hz) (Faivre et al., 2018; Samaha et al., 2017; Wöstmann et al., 2019). However, these EEG signatures have rarely been examined outside the domain of perception and hence the degree to which they relate to either domain-specific or domain-general mechanisms is unknown.

In addition to debate regarding domain-generality, the test-retest reliability of both behavioural and neurophysiological measures of metacognition also remains unclear. Though computational models of behaviour, like those often employed to estimate metacognitive performance (Fleming, 2017; Maniscalco & Lau, 2012), offer insights into latent processes, increasing concerns have been raised about their reliability and potential to capture trait-like characteristics (Brown et al., 2020; Hedge et al., 2018; Shahar et al., 2019). Likewise, the reliability of neuroimaging based bio-markers of cognitive functions have also been called into question (Botvinik-Nezer et al., 2019; Botvinik-Nezer & Wager, 2022; Haines et al., 2023; Pavlov et al., 2021; Poldrack et al., 2017).

We examined the relationships between both pre- and post-stimulus EEG activity and subjective confidence across separate tasks that engage distinct cognitive processes – visual perception and semantic memory. Additionally, we investigated the test-retest reliabilities of both behavioural and neural measures of metacognition across two experimental sessions. We found that overall confidence calibration (i.e., metacognitive bias) was the most reliable measure across cognitive domains and time, whereas metacognitive efficiency showed poor test-retest reliability and low domain-generality. Fluctuations of confidence within-participants were reliably captured by stimulus-locked event-related potential (ERP) activity across cognitive domains, and less reliably captured by event-related spectral perturbation (ERSP) activity.

## 2. Materials and Methods

### 2.1. Participants

We recruited twenty-seven participants, of whom twenty-five completed two experimental sessions on two separate days. Only the 25 who completed two sessions were included in the analyses (15 females, 24 right-handed, 1 ambidextrous, age: *M* = 23.2, *SD* = 3.4, range = 18-34). The mean number of days between testing sessions was 25 (range = 2-85). Inclusion criteria were age between 18 and 40 years, normal or corrected-to-normal vision, and no history of neurological or psychiatric disorders. Ten participants wore glasses or contact lenses. On average, participants reported sleeping *M* = 7.21 (*SD* = 1.43) hours the night prior to testing. All participants gave written informed consent in accordance with the Declaration of Helsinki and were financially compensated for their time (£30 per recording session). The study was approved by the School of Social Sciences Ethics Committee at the University of Dundee and took place within the University’s Psychology department.

### 2.2. Tasks and Experimental design

Participants completed two separate 2-alternative forced-choice tasks, one perceptual (Benwell et al., 2022b; Rouault, Seow, et al., 2018) and one semantic knowledge based (Benwell et al., 2022b; Sanders et al., 2016), in a counterbalanced (blocked) order whilst 64-channel EEG was simultaneously recorded. Each participant completed two experimental sessions on separate days thereby allowing us to examine the test-retest reliability of both the behavioural and EEG results. The order of the tasks was counterbalanced between the experimental sessions. The perceptual task (PT) involved deciding which of two simultaneously presented boxes presented contained a higher number of dots, while in the knowledge task (KT) participants were required to choose which of two countries had a larger human population. Following each response, confidence judgements were provided on a six-point rating scale (see Figure 1).

**Figure 1:**
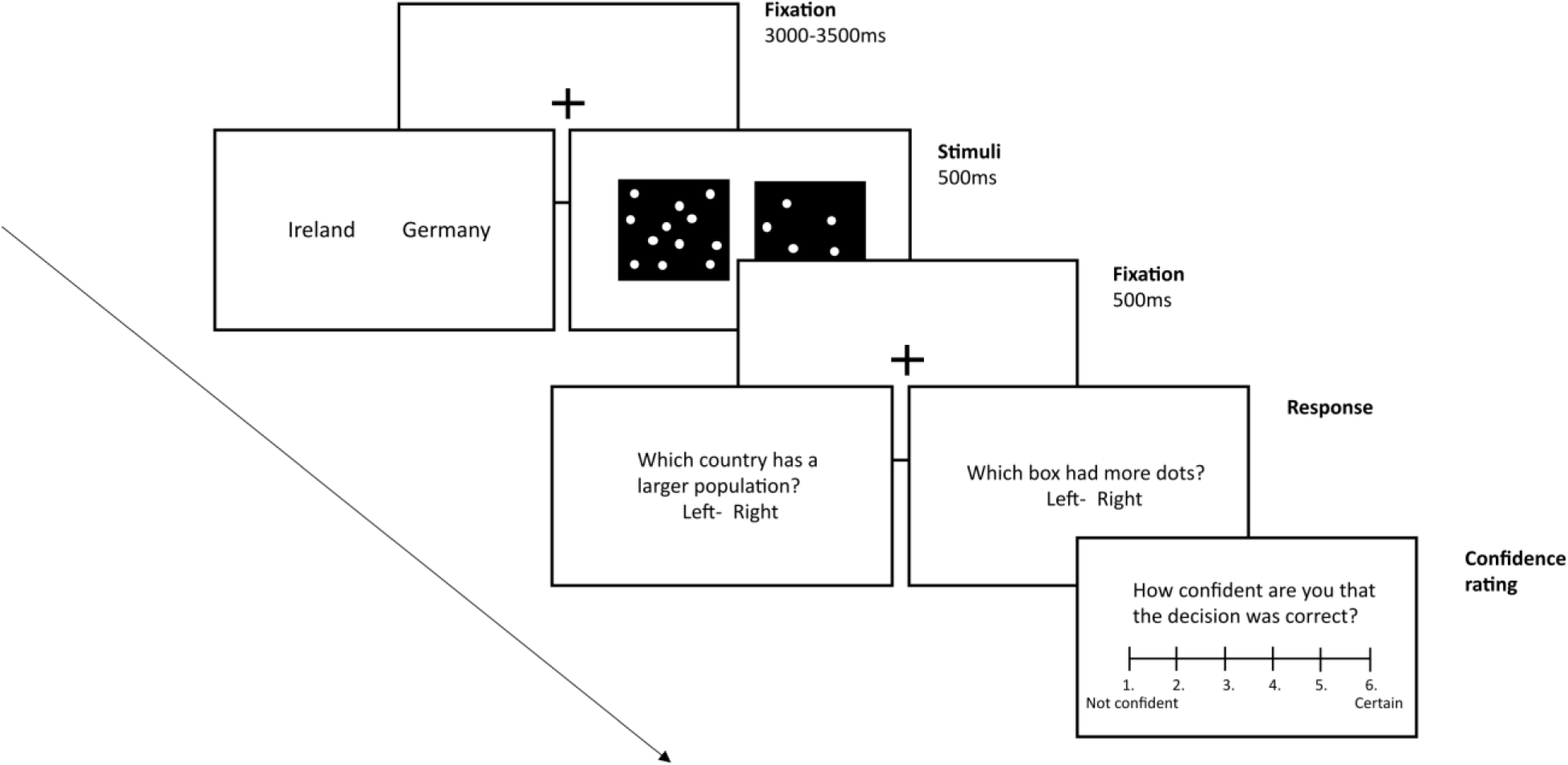
Perceptual and knowledge tasks. Both experimental tasks started with a black fixation cross presented on a white background for a randomly varying duration of 3000-3500ms. Experimental stimuli followed, remaining on the screen for 500ms and consisting of pairs of country names for the *knowledge task* and pairs of black boxes with varied number of white dots for the *perceptual task*. Next, another fixation cross appeared for 500ms, followed by the response prompt then the confidence rating scale, which remained on the screen until participants responded.

We presented all stimuli using PsychoPy software (Peirce et al., 2019) on a 53x30cm monitor with 1920x1080 resolution and 60Hz refresh rate. Participants sat in a dimly lit room approximately 70cm from the computer screen. Each trial started with a black fixation cross (1.5x1cm) presented on a white background for a duration randomly varying between 3000-3500ms followed by the experimental stimuli presented for 500ms. For the perceptual task, stimuli consisted of two black boxes (8.5x9cm) containing white dots, one on the left and the other on the right side of the screen (with 4.2cm between the boxes). One box (the reference box) constantly contained 272 dots (out of 544 possible dot locations), while the other box contained an increased or reduced number of dots ranging from either −40 to +40 dots (in increments of 8) in comparison to the reference box (including an identical condition). For each difficulty level, the position of the target box (left/right) was randomised. For the knowledge task, stimuli consisted of two country names whose population ratios were varied across 5 difficulty levels. We downloaded the national populations from The World Bank (‘https://data.worldbank.org/indicator/SP.POP.TOTL’) in December 2019. We created five different evidence discriminability ‘bins’ by grouping country pairs with similar population log ratios (log10(Country-A Population/Country-B Population)). The log ratio bins amounted to the following, ranging from least to most discriminable: bin 1 (log10 ratio = 0-0.225), bin 2 (0.225-0.45), bin 3 (0.45-0.675), bin 4 (0.675-0.9), bin 5 (0.9-1.125). Each bin included 18 different country pairs. The left/right position of names in each country pair was randomised across participants. The stimuli were followed by another fixation cross that lasted 500ms. The inclusion of this time-period prior to response ensured that stimulus induced EEG waveforms were not contaminated by motor activity. Afterwards, participants were prompted to give their response using the ‘w’ (left larger) and ‘e’ (right larger) keys. Next, they reported their decision confidence by pressing one of six keys on the numeric pad (1-not confident at all, 6-certain). The response and confidence prompts remained on the screen until participants responded. To familiarise themselves with the tasks, participants completed ten practice trials prior to each. In total, each experimental session consisted of 400 (knowledge) and 440 (perceptual) task trials, each split into 5 blocks, with trial order fully randomised within blocks. This resulted in 80 trials per difficulty level in each task, with forty additional catch trials included in the perceptual task. Overall, each recording session lasted approximately 2.5-3 hours including EEG set-up. We randomised the order of task performance across participants.

### 2.3. Behavioural analysis

#### Model-free analyses

To evaluate the effectiveness of the task difficulty manipulation on each task, we compared the mean accuracy and confidence ratings across all difficulty levels using one-way repeated-measures Analyses of Variance (ANOVA), with difficulty as the independent variable and proportion of correct responses/mean confidence as the dependent variables. Where Mauchly’s Test of Sphericity indicated unequal variances (*p* < .05), we applied the Greenhouse-Geisser correction (𝑝_𝐺𝐺_). We used Eta squared (*η*^2^) as measure of effect sizes.

#### Modelling metacognition

We modelled 1^st^-order decisions and subjective confidence ratings in both tasks within an extended signal detection theory (SDT) framework (Maniscalco & Lau, 2012). All parameters were modelled using data collapsed across all difficulty levels. We calculated both type-1 sensitivity (***d’***, indexing 1^st^-order decision accuracy) and type-2 (metacognitive) sensitivity (***meta-d’***, equal to the value of ***d*’** required to produce the observed confidence ratings data by an optimal metacognitive observer with the same type-1 criterion) for each participant. A metacognitively optimal observer should have ***d’*** equal to ***meta-d’***. We then calculated the metacognitive ratio (***meta-d’/d’***) as a measure of metacognitive efficiency (estimate of metacognitive sensitivity whilst controlling for 1^st^-order performance), whereby *M-ratio* = 1 suggests ideal efficiency, *M-ratio* < 1 suggests not all evidence available for type-1 decisions was used to make type-2 decisions, and *M-ratio* > 1 suggests more evidence was available for type-2 than type-1 decisions (Fleming & Lau, 2014). A measure of confidence bias, the type-2 (confidence) criterion (***type-2 c’***), was also calculated to estimate participants tendency to give high/low confidence ratings regardless of overall accuracy. To separate confidence bias from type-1 response bias, we computed the absolute difference between the type-1 and type-2 criteria (Benwell et al., 2022b; Sherman et al., 2018). We averaged the type-2 criterion across N-1 available confidence ratings and then across both possible responses (‘left’/’right’) for all analyses. All of the measures were estimated using the Maximum Likelihood Estimation method from Maniscalco & Lau, (2012) (http://www.columbia.edu/~bsm2105/type2sdt/) using the ‘fit_meta_d_MLE’ function in MATLAB R2021a (Mathworks, USA).

#### Test-retest reliability & domain-generality

To test the reliability of all behavioural measures, both model-free (accuracy, confidence) and model-based, we calculated Intraclass correlation (ICC) coefficients both between experimental sessions and between tasks. Two-way consistency ICC type was used (C-1, based on the McGraw and Wong (1996) convention) and calculated with the ICC function in MATLAB (Salarian, 2023 (‘https://www.mathworks.com/matlabcentral/fileexchange/22099-intraclass-correlation-coefficient-icc’)). We also used paired-samples t-tests to test for between-task differences on each day for each measure of interest.

### 2.4. Electroencephalography acquisition and pre-processing

Continuous EEG was recorded using a 64-channel ActiveTwo system (Biosemi, The Netherlands) at a sampling rate of 1024Hz. The scalp electrodes were placed according to the International 10-20 system. We placed four additional electrooculographic electrodes at the outer canthi of each eye as well as above and below the participant’s right eye to capture horizontal and vertical eye movements respectively.

We performed EEG data pre-processing offline using custom-written scripts in MATLAB R2021a (Mathworks, USA) including EEGLAB (Delorme & Makeig, 2004) and Fieldtrip (Oostenveld et al., 2011) functions. Low-pass (100Hz) and high-pass (0.5Hz) filters were applied to the data using a zero-phase second-order Butterworth filter. We then divided the filtered recordings into 4 second epochs, from −2.5 to 1.5s relative to stimulus onset on each trial. We visually inspected the data and removed faulty or excessively noisy channels without interpolation (Day 1: range=0-3, *M*=0.4 (KT), 0.44 (PT); Day 2: range 0-3, M = 0.56 (KT), M = 0.36 (PT)). The recording was subsequently re-referenced to the average of all channels (excluding the four non-scalp electrodes) and excessively noisy trials were removed following a semi-automated artifact identification procedure in which trials with potential artefacts were identified based on (1) extreme amplitudes (threshold of ± 75 µV), (2) joint probability of the recorded activity across electrodes at each time point (probability threshold limit of 3.5 and 3 standard deviations [SD] for single-channel limit and global limit, respectively; pop_jointprob; Delorme & Makeig, 2004) and (3) kurtosis (local limit of 5 SD, global limit of 3 SD; pop_rejkurt; Delorme & Makeig, 2004). We then ran independent component analysis (ICA) using the ‘*runica’* function in EEGLAB (Delorme & Makeig, 2004) and components corresponding to blinks/eye-movements, muscle activity, or transient channel noise were semi-automatically removed using the Multiple Artifact Rejection Algorithm (MARA) (Winkler et al., 2011). Next, we removed any remaining noisy epochs, this resulted in a mean of 424 (range = 401-437) perceptual and 386 (range = 369-398) knowledge trials per participant (across both testing sessions). We excluded the rejected trials from all behavioural analyses too. Finally, previously removed channels were interpolated using a spherical spline method.

For ERP analyses only, we applied an additional low-pass (40Hz) filter to the clean time-series data. The data were then cut into epochs spanning −500:1000ms relative to stimulus onset on each trial and baseline corrected using the 500ms pre-stimulus period.

#### Time-frequency spectral power analysis

To estimate spectral power across time and frequency domains, we performed a Fourier-based time-frequency transformation on the clean single-trial data for each channel using the ‘*ft_freqanalysis*’ function (method: ‘*mtmconvol*’) in the Fieldtrip toolbox (Oostenveld et al., 2011). Overlapping sections of single-trial time-series data were decomposed by consecutively shifting a 0.5-second-long window forward in time by 0.02s and Hanning tapered. A frequency resolution of 1Hz across 1-40Hz range was achieved by zero-padding the data to a length of 1 second. The absolute values of the resulting complex-valued time-frequency estimates were then squared to obtain single-trial spectral power and converted into decibels (dB). We then used data epochs spanning −1:1s relative to stimulus onset in the statistical analyses.

### 2.5. Statistical analyses

We carried out all statistical analyses in MATLAB R2021a and 2022a (Mathworks, USA) using custom-written code.

#### Single-trial regression analysis

To elucidate both the ERP and pre- and post-stimulus time-frequency spectral power correlates of subjective confidence ratings, we adopted a non-parametric single-trial regression approach (Benwell et al., 2017, 2022a; Samaha et al., 2017). The large number of task trials per participant allowed us to test for systematic EEG-behaviour relationships in a hierarchical two-stage estimation model (Friston, 2008), thus incorporating subject-level variation into the group-level statistics. Prior to the regression analyses, we rank transformed all variables of interest apart from accuracy to nullify the influence of any outlying trials.

First, we tested *within-participant relationships* between spectral power and confidence ratings with separate regression models for each univariate response (i.e., time-electrode point (for ERPS) or time-frequency-electrode point (for time-frequency power)). The single-trial EEG power (𝐸𝐸𝐺), accuracy (Accuracy), and difficulty level (Difficulty) were entered as predictors of the behavioural (Confidence) outcome. This allowed us to test the relationships between EEG activity and confidence whilst controlling for external evidence strength and 1^st^-order accuracy.

We obtained the regression coefficients for the EEG-confidence relationship from the following multiple linear regression model:

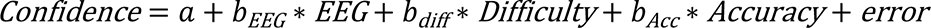

where 𝑏_𝐸𝐸𝐺_ represents the strength and direction of the unique relationship between EEG activity and confidence. The linear regression models were applied using the ‘*fitlm’* function with the least-squares criterion (Mathworks, USA).

Second, we computed *group-level* statistics with cluster-based permutation testing (Maris & Oostenveld, 2007) using the *‘ft_freqstatistics’* function in Fieldtrip (method: ‘*montecarlo’*, Oostenveld et al. 2011). Regression coefficients (𝑏_𝐸𝐸𝐺_) at each data point (time-frequency-electrode) were combined across all participants, and a two-tailed dependent-samples t-test against zero was used to test for systematic group-level effects. We used cluster-based permutation testing to control the family-wise error rate (Maris & Oostenveld, 2007). First, clusters were formed by combining all adjacent significant time-frequency-electrode datapoints based on the initial t-tests, separately for negative and positive values, and summing the t-values to produce a cluster-level statistic. A minimum of 1 significant neighbouring sample was required to form a cluster. Electrodes were considered as neighbours based on the *‘biosemi64-neighb.mat*’ template in Fieldtrip (Oostenveld et al., 2011), created through symmetric triangulation and manual editing, and leading to a mean of 6.6 (range=3-8) neighbours per channel. The cluster t-statistics were then compared against a data-driven null hypothesis distribution. This was obtained by randomly drawing coefficients from a subset of participants, multiplying them by −1, thereby cutting the hypothesised brain-behaviour relationship, and forming clusters based on significant t-tests against 0 as described above. We repeated the whole procedure 2000 times while saving the largest cluster t-statistic on each iteration, thus building a distribution of t-values that would be expected in the absence of an EEG-behaviour relationship. Consequently, the t-statistics of the true positive and/or negative clusters falling within either the lower or higher 2.5% of the null distribution (alpha = .05, two-tailed) were considered significant.

#### Decoding analysis

Finally, to investigate the domain-generality and test-retest reliability of confidence-related EEG activity, we performed a series of multivariate decoding analyses. First, within each task on each day separately, we tested whether single-trial confidence could be decoded from post-stimulus EEG activity (both ERP and spectral power respectively). Crucially, we then performed a cross-decoding analysis which allowed us to investigate whether a classifier trained on a task from one cognitive domain could also predict confidence in the other cognitive domain. A significant cross-classification would indicate that shared neural representations underlie confidence judgements across cognitive domains, providing evidence for domain-general neural correlates of metacognition. If cross-classification is not possible or is weaker than within-task classification, perceptual and knowledge decision confidence may be related to domain-specific or only partially overlapping neural mechanisms. Similarly, we also investigated the test-retest reliability of the within-task decoding by testing whether a classifier trained on the data from the first day could decode confidence on the second day as well.

We performed all multivariate pattern analyses using the MVPA Light toolbox (Treder, 2020) in MATLAB 2022a (Mathworks, USA). Classifiers were trained to decode between high and low confidence trials which we determined using a binning method that ensured the most even split between high/low confidence trials within each participant. We calculated the number of trials for each of the 6 possible confidence ratings and then binned the trials so that the difference in trial number between high and low confidence bin was minimised (mean difference day 1 = 15.52% in a range 0.25-42.65%; day 2 = 14.64% in a range 1.52-44.25%). This procedure is equivalent to a within-participant median split of continuous data. Across datasets from both days (n = 50), 5 datasets were in a bin split into [1 = low bin] [2 3 4 5 6 = high bin]; 5 in a split of [1 2] [3 4 5 6]; 10 in [1 2 3] [4 5 6]; 12 in [1 2 3 4] [5 6]; and 18 in [1 2 3 4 5] [6].

We used a Linear Discriminant Analysis (LDA) classifier and calculated the area under the receiver operating characteristic curve (AUC) as a classifier performance metric. For within-task decoding, we used 5-fold cross-validation to avoid overfitting and repeated it 10 times to reduce the noise following the random assignment of trials into each fold. For analysis of ERP data, we computed a classifier that identifies the spatial distribution of EEG activity (across all 64 electrodes) which maximally distinguished between high and low confidence at each time point (0:1s). To assess the temporal generalisation of the classification performance, all classifiers were trained and tested on each time point of the data, thereby producing a 2D matrix of decoding performance (train x test time). The shape of the final decoding matrix allows inferences about the temporal dynamics of mental representations (King & Dehaene, 2014). Similarly, for the analysis of spectral power, we performed a classification analysis using the average power in the classic alpha (8-12Hz) and beta (13-30Hz) bands across all post-stimulus time points (0:1s), training and testing the classifier at each time point. We trained the classifiers using the average power from individual bands as well as their combination.

We performed the cross-task and cross-day decoding using an LDA classifier with AUC performance metric as well. In the between-task classification, we first trained the classifier on the whole dataset (combined across both testing days) from the knowledge task and then tested it on data from the perceptual task (and vice versa). We examined test-retest reliability by training the classifier on day 1 data and testing it on day 2 data for each task separately.

The statistical significance of all classifiers at the group level was tested using cluster-based non-parametric permutation testing (Maris & Oostenveld, 2007) similar to that we employed in the single-trial regression analyses. We compared the actual classification values at all datapoints (in a 2D matrix for temporal generalisation) to a null classification AUC value (0.5) with a paired-samples t-test. An element was considered significant if *p* < .05 (one-tailed). All neighbouring datapoints that passed the element-level significance threshold were collected into a cluster. Directly and diagonally adjacent elements were considered as neighbours. We obtained a cluster-level statistic by summing the individual t-values within each cluster. We then compared this against a null distribution generated by repeating the whole procedure 2000 times while randomly permuting the data. Clusters with cluster-level *p* < .05 were considered significant.

## 3. Results

### 3.1 Behavioural results

First, we evaluated and compared the overall performance on both the perceptual and knowledge tasks. Figure 2A (day 1) and Figure 2B (day 2) plot the group-averaged proportion of correct responses as a function of evidence discriminability for both tasks. As expected, in both tasks (and on both days), the proportion of correct responses increased significantly from the hardest to the easiest trials (Perceptual task day 1: F(1,24) = 151.71, *p_GG_*<.0001, *η*^2^= 0.863, all post-hoc *p*’s <.05, Perceptual task day 2: F(1,24) = 153.77, *p_GG_*<.0001, *η*^2^= 0.865, all post-hoc *p*’s <.05 apart from bins 4-5 (*p* = .734); Knowledge task day 1: F(1,24) = 247.55, *p* <.0001, *η*^2^= 0.912, all post-hoc *p*’s <.05, Knowledge task day 2: F(1,24) = 161.99, *p* < .0001, *η*^2^= 0.871, all post-hoc *p*’s < .05). Comparisons of proportion of correct responses between the tasks, performed separately at each evidence discriminability level, showed that on both days participants were significantly more accurate in the perceptual task at all difficulty levels apart from the easiest condition (all *p*’s < .05). The scatterplots in Figure 2A-B further show that the overall proportion of correct responses on the perceptual and knowledge tasks were not significantly correlated in either experimental session (ICCs day 1: r = .287 *p* =.078; day 2: r = .289 *p* =.076). Hence, 1^st^-order accuracy did not show strong domain-generality across tasks. However, it did show strong test-retest reliability from session 1 to session 2 within both tasks (Figure 2C: perception r = .817, *p* < .0001; knowledge r = .807, *p* < .0001). Figure 2C also shows that overall perceptual task accuracy increased from day 1 (mean proportion correct: M = 0.824) to 2 (M = 0.842) (t(24) = −2.227, *p* = .036, Cohen’s d = 445), whereas overall knowledge task accuracy did not significantly differ from day 1 (M = 0.753) to 2 (M = 0.743) (t(24) = 1.157, *p* = .259, d = 231).

**Figure 2:**
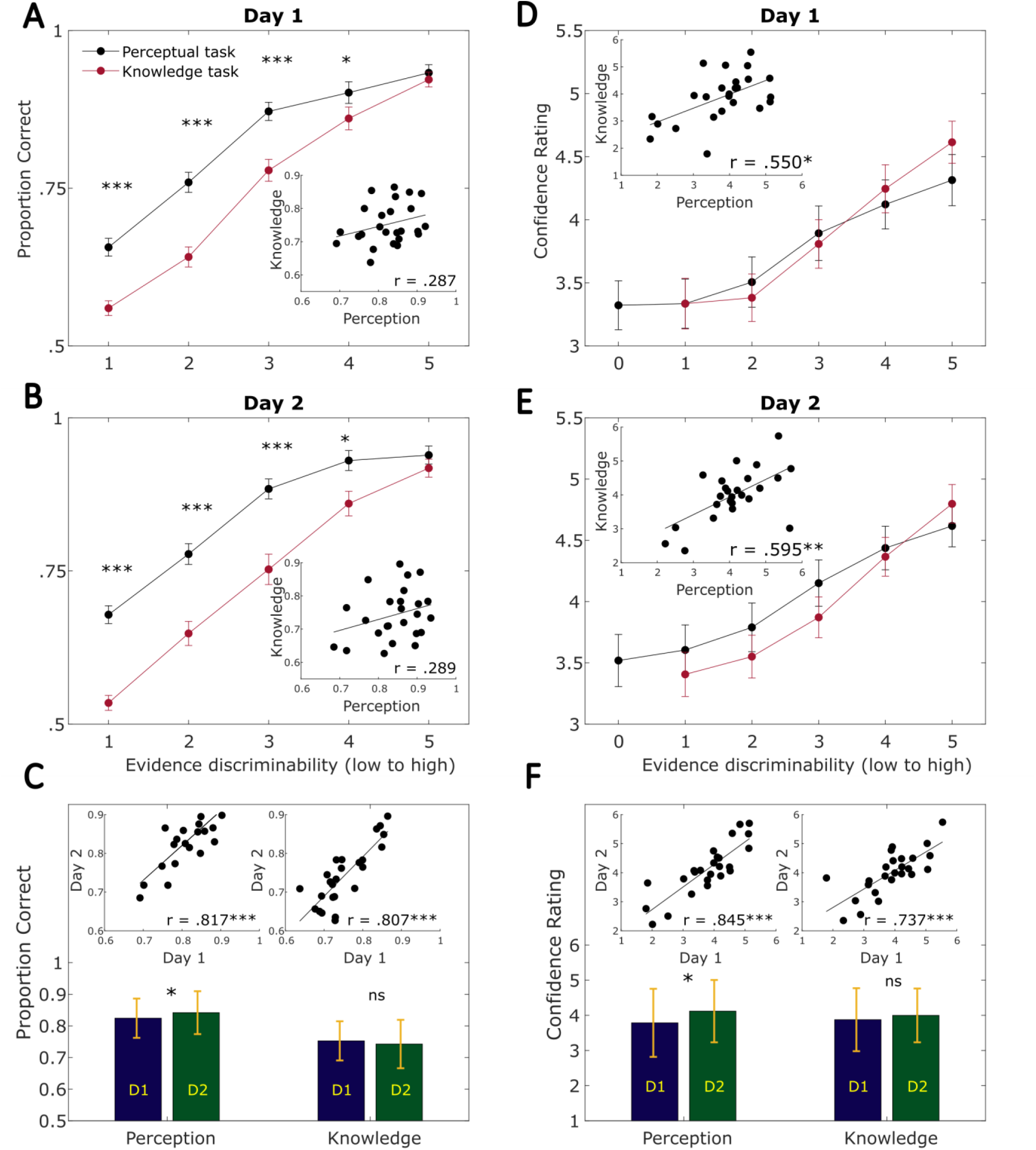
Model-free behavioural performance. **A-B** Group-averaged proportion of correct responses per evidence discriminability level on each testing day. Error bars represent between-participant standard error of the mean (SEM). Scatterplots show the relationship between the proportion correct responses from each task (with Intraclass correlation coefficients (two-way consistency type)). **C** Comparisons of objective performance between testing days. The bar chart shows mean proportion correct for each task on each day. Error bars represent standard deviation. The scatterplots show the test-retest reliability of objective performance in each task separately, measured by intraclass correlations (ICCs). **D-E** Group-averaged confidence ratings per difficulty level. Error bars represent SEM. Scatterplots show the correlation in confidence rating between tasks. **F** Test-retest reliability (scatterplots) and between-day comparison of confidence ratings (bar chart, error bars = SD) in each task. *** *p* < .0001, ** *p* < .001, * *p* < .05

Next, we investigated subjective confidence across tasks. Figure 2D (day 1) and Figure 2E (day 2) plot group-averaged confidence ratings as a function of evidence discriminability for both tasks. As expected, average confidence ratings increased significantly as task difficulty decreased (Perceptual task day 1: F(1,24) = 50.36, *p_GG_*<.0001, *η*^2^= 0.677, all post-hoc *p*’s < .05 apart from comparison in bins 0-1 (*p* =.9995), 0-2 (*p* = .091), Perceptual task day 2: F(1,24) = 35.21, *p_GG_*<.0001, *η*^2^= 0.595, all post-hoc *p*’s < .05 apart from bins 0-1 (*p* = .491), Knowledge task day 1: F(1,24) = 72.71, *p_GG_*<.0001, *η*^2^= 0.752, all post-hoc *p*’s < .05 apart from bins 1-2 (*p* =.678), Knowledge task day 2: F(1,24) = 57.00, *p_GG_*<.0001, *η*^2^= 0.704, all post-hoc *p*’s < 05 apart from bins 1-2 (*p* =.067). Despite 1^st^-order accuracy being higher for the perceptual task across most difficulty levels (Fig 2A-B), we did not observe any significant between-task differences in average confidence at any of the individual difficulty levels on either of the testing days (Fig 2D-E: all *p*’s > .05). In contrast to 1^st^-order accuracy, overall average confidence ratings were strongly correlated between the tasks on both testing days (Figure 2D-E scatterplots; ICCs day 1: r = .550 *p* =.0018, day 2: r = .595 *p* =.0007). Hence, unlike accuracy, confidence ratings showed reliable and moderately strong domain-generality across the cognitive domains tested (perception and semantic memory). Average confidence ratings also showed excellent test-retest reliability from session 1 to session 2 within both tasks (Figure 2F: ICCs perception r = .845, *p* < .0001; knowledge r = .737, *p* < .0001). In line with 1^st^-order accuracy, Figure 2F shows that overall confidence increased from day 1 (M = 3.787) to day 2 (M = 4.120) (t(24) = −3.222, *p* = .004, d = .645) for the perceptual task, but not for the knowledge task (t(24) = −1.012, *p* = .322; day 1 M = 3.877, day 2 M = 3.999, d = .202).

Overall, in both tasks the difficulty manipulation was effective, and participants used the confidence scale appropriately. Test-retest reliabilities of both confidence and objective accuracy were high within-tasks, but only confidence showed a significant between-task correlation indicating domain-generality.

#### Model-based metacognition

We next investigated the reliability and domain-generality of model-based measures of performance (collapsed across all discriminability levels) derived from an extended signal detection theory (SDT) framework (Maniscalco & Lau, 2012). Comparison of type-1 and type-2 performance measures between the tasks (Figure 3) showed that type-1 sensitivity (***d’***) was significantly higher in perception than knowledge on both days (D1: perception M(SD) = 1.806(0.443), knowledge M(SD) = 1.392(0.416), t(24) = 4.118, *p* < .001, Cohen’s d = .824; D2: perception M(SD) = 2.281(0.558), knowledge M(SD) = 1.344(0.509), t(24) = 7.257, *p* < .0001, d = 1.451), in line with the proportion correct results above. Overall ***type-2 (confidence) c’*** (indexing confidence bias) did not differ significantly between-tasks on either day (D1: perception M(SD) = 0.959(0.561), knowledge M(SD) = 0.831(0.452), t(24) = 1.407, *p* = .172, d = .281; D2: perception M(SD) = 0.891(0.492), knowledge M(SD) = 0.758(0.339), t(24) = 2.047, *p* = .052, d = .409), whereas metacognitive efficiency (***Meta-d’/d’***) was significantly higher for knowledge compared to perception (D1: perception M(SD) = 0.709(0.225), knowledge M(SD) = 0.927(0.259), t(24) = −3.914, *p* < .001, d = .783; D2: perception M(SD) = 0.675(0.244), knowledge M(SD) = 0.990(0.209), t(24) = −5.754, *p* < .0001, d = 1.151). Hence, in line with our previous study (Benwell et al., 2022b), whilst 1^st^-order performance was significantly better on the perception task, metacognitive efficiency was significantly higher for the knowledge task. Higher metacognitive efficiency in the knowledge task may be due to differences in evidence types between tasks. Unlike perceptual evidence, semantic evidence in the knowledge task is internal and continuously available throughout the trial for metacognitive evaluation.

**Figure 3:**
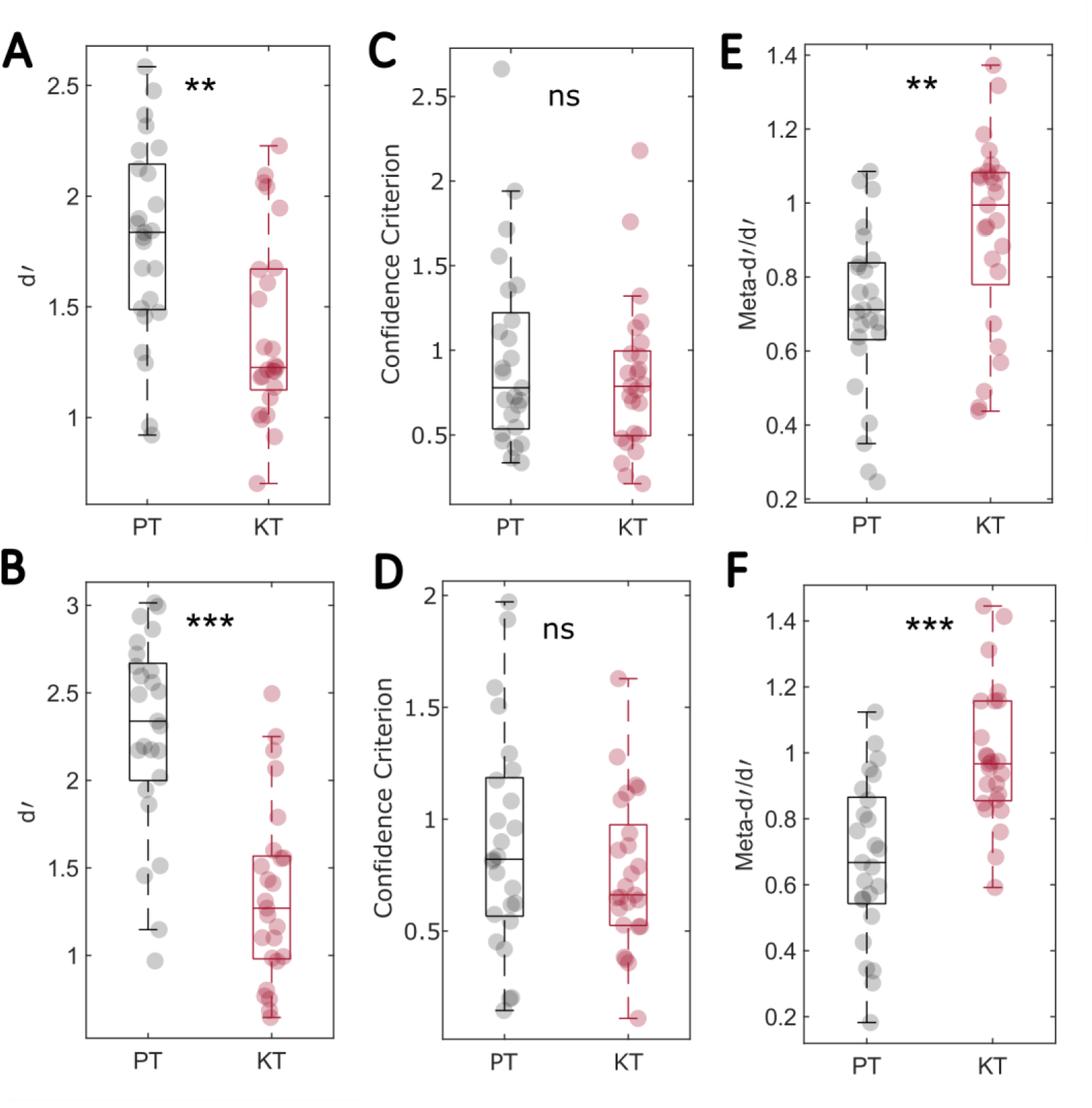
Between-task comparisons of model-based type1 and type-2 performance measures. The top row plots day 1 data and the bottom row plots day 2 data. **A-B** type-1 sensitivity (d’), **C-D** type-2 (confidence) criterion, and **E-F** metacognitive efficiency (M-ratio). The central boxplot lines correspond to the median, the upper box edges correspond to the 0.75 quantile and the lower edges represent the 0.25 quantile. The whiskers represent the non-outlier maximum and minimum values.

To test the domain-generality of the model-based metacognitive measures, we computed between-task intra-class correlations for each. We reasoned that significant correlation of a given measure between tasks suggests that a shared mechanism must contribute across cognitive domains. As Table 1 shows (see Supplementary Figure S1 for scatterplots), the only measure that showed a strong and replicable significant between-task correlation was ***type-2 (confidence) c’***, with r = .601 on day 1, and .705 on day 2. This suggests that overall confidence calibration (metacognitive bias) strongly contributes to metacognition in a domain-general manner. In contrast, neither type-1 sensitivity (***d’***) nor metacognitive efficiency (***Meta-d’/d’***) showed replicable significant between-task correlations. Although metacognitive efficiency showed a weak significant correlation between tasks on day 1, this was not replicated on day 2.

**Table 1:**
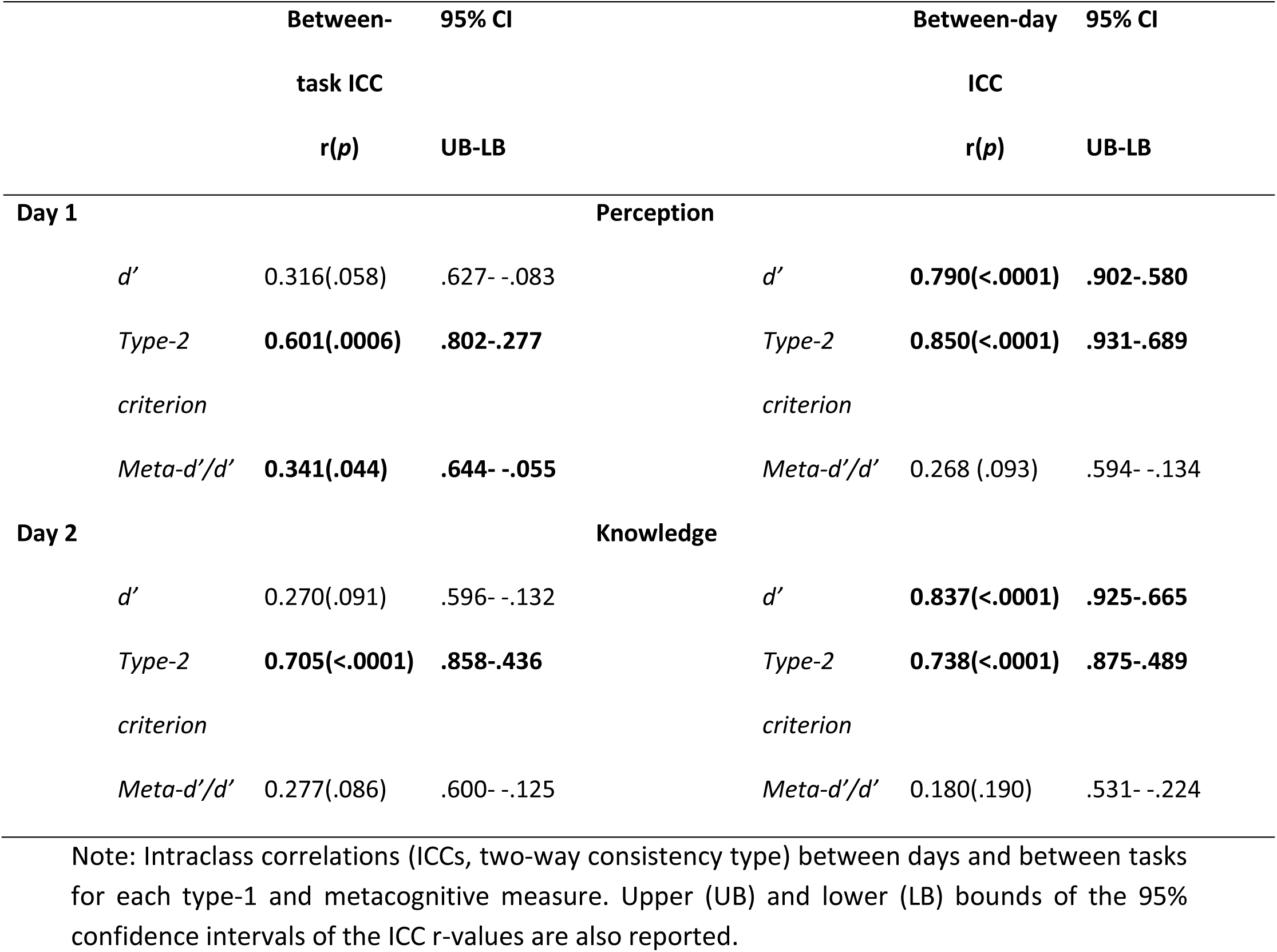
Domain-generality and test-retest reliability of Type-1 and Type-2 performance

We tested the test-retest reliability of all measures with between-day, within-measure intra-class correlations (Table 1, see also Supplementary Figure S2). Both ***d’*** and ***type-2 c’*** were strongly correlated across both testing sessions for both tasks (all p’s < .0001, r range = .738-.850) suggesting excellent test-retest reliability of type-1 sensitivity and confidence bias. In contrast, metacognitive efficiency (***Meta-d’/d’***) did not significantly correlate between testing sessions for either task, indicating poor test-retest reliability of this measure.

Summarising the results for our metacognitive behavioural measures, overall confidence calibration (as indexed by both average confidence ratings and ***type-2 c’***) was both domain-general and highly reliable over time, whereas metacognitive efficiency (as indexed by ***Meta-d’/d’***) was neither domain-general nor reliable over time. Hence, we focussed our EEG analyses on identifying reliable signatures associated with single-trial confidence reports across cognitive domains.

### 3.2 EEG results

To identify EEG predictors of confidence during decision formation, we investigated single-trial relationships between confidence ratings and both ERP and time-frequency spectral power activity for each task separately across both experimental sessions.

#### Late stimulus-locked ERP activity reliably predicts domain-general subjective confidence

We used cluster-based permutation analysis (Maris & Oostenveld, 2007) to identify systematic relationships between single-trial ERP amplitude and confidence ratings across all electrodes and post-stimulus time points (0-1s relative to stimulus presentation). Whilst controlling for difficulty level and 1^st^-order accuracy, we observed significant relationships between a late component of the evoked potential and confidence ratings in both tasks which replicated across both experimental sessions (Figure 4).

**Figure 4:**
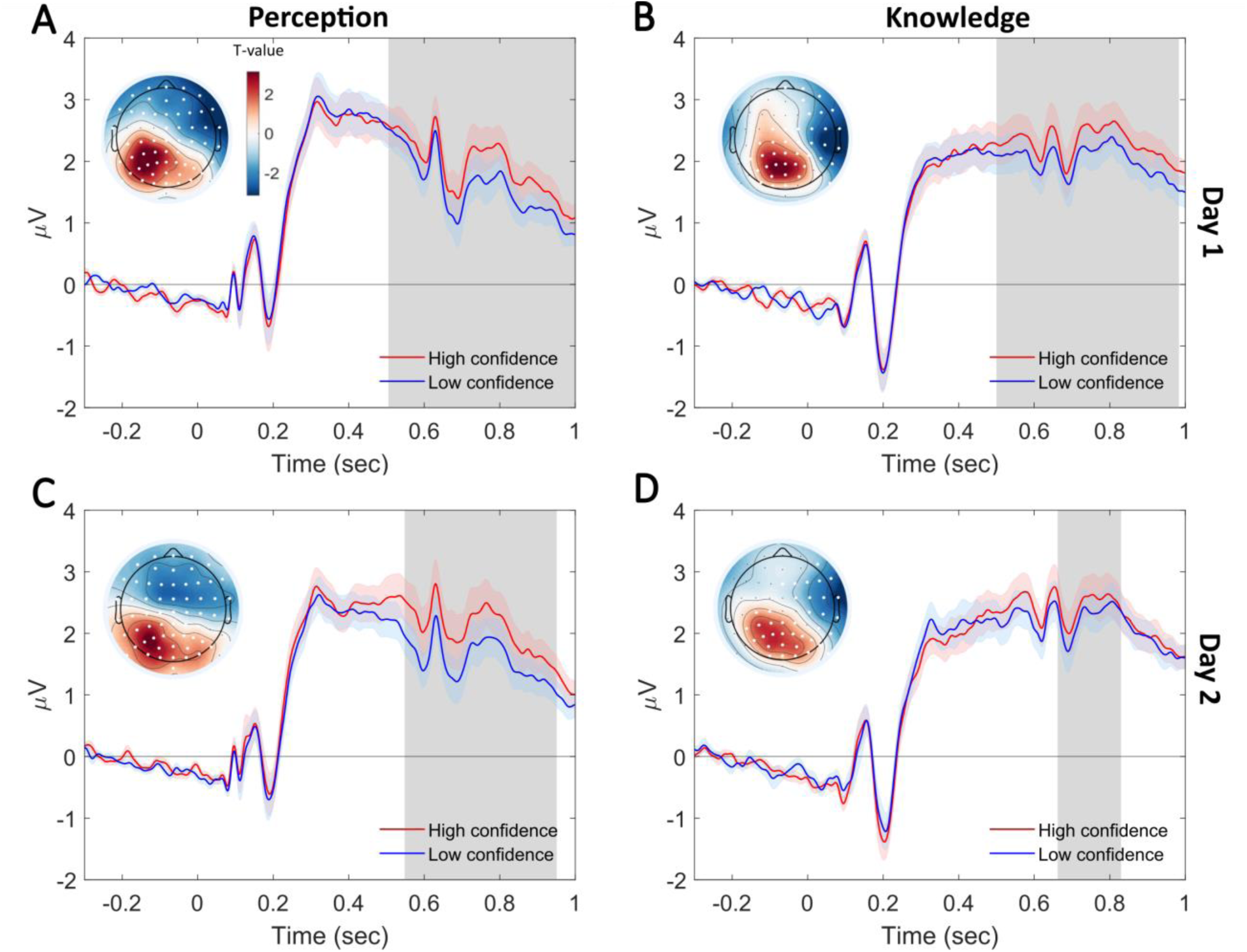
Late ERP activity reliably reflects subjective confidence ratings, independently of accuracy and evidence discriminability, across cognitive domains. Grand-averaged stimulus-locked ERP waveforms at electrode P3 for high (red) and low (blue) confidence trials are shown for the **A** Perception task (day 1), **B** Knowledge task (day 1), **C** Perception task (day 2), and **D** Knowledge task (day 2). The high versus low confidence trials were defined in a binning procedure that minimised the difference in the number of high/low confidence trials within each participant (i.e., approximate median split of ordinal data, see Methods). Red/blue shaded waveform areas represent SEM of the respective waveforms. Note that we only performed the binning here to plot high and low confidence waveforms for illustrative purposes. We performed the statistical analysis on single-trial (non-binned) EEG and behavioural data. Gray shaded areas represent the significant time windows for positive clusters, as identified through cluster-based permutation testing. The topographies plot the mean t-values for all time-points from the significant cluster across participants, with both significant positive cluster electrodes and negative cluster electrodes highlighted in white.

For the perceptual task on day 1, we found that two significant clusters predicted confidence (Figure 4A): one positive (cluster statistic = 17,693, *p* = .0015; spanning ∼490-1000ms post-stimulus over centroparietal/occipital electrodes with a left parietal maximum (see topography in Figure 4A)) and one negative (cluster statistic = −18,845, *p* = .001; spanning ∼520-1000ms post-stimulus over frontal electrodes with a right maximum). We found two highly similar clusters that predicted confidence on the knowledge task on day 1 as well (Figure 4B): one positive (cluster statistic = 11,124, *p* = .0025; ∼490-960ms over centroparietal electrodes (see Figure 4B topography)) and one negative (cluster statistic = −9,953 *p*= .0015; ∼470-890ms over right frontotemporal electrodes).

These results were largely replicated in the second testing session (Figure 4C-D). On day two, we found a positive (cluster statistic = 14,267, p = .002, beginning at ∼530ms, over left centroparietal/occipital electrodes, Fig. 4C) and a negative (cluster statistic = −17,997, p = .0005, starting at ∼530ms, over right frontotemporal electrodes) cluster in the perceptual task. In the knowledge task (day 2, Fig. 4D), we found a positive cluster (cluster statistic = 4,360, p = .0265, spanning ∼640-810ms, over left centroparietal/occipital electrodes) and a negative cluster (cluster statistic = −6,387, p = .0125, spanning ∼480-930ms, over right frontotemporal electrodes).

Based on the topography and timing of the observed clusters, they possibly represent opposite dipoles of the late portion of the P3/central parietal positivity (CPP) component previously implicated in decision-making and subjective measures of perception including confidence (Azizi et al., 2021; Feuerriegel et al., 2022; Fitzgerald et al., 2022; Gherman & Philiastides, 2015, 2018; Herding et al., 2019; Lim et al., 2020; Tagliabue et al., 2019). Participants’ confidence tended to be higher on trials with stronger late CPP amplitude irrespective of the cognitive domain tested, thus highlighting that the CPP reliably reflects domain-general subjective confidence over and above the external stimulus information (i.e., difficulty) and objective accuracy.

#### Post-stimulus, but not pre-stimulus, alpha/beta desynchronisation predicts subjective confidence

We additionally tested whether single-trial spectral power across time (−1s:1s) and frequencies (1:40 Hz) is related to confidence ratings. We included the 1s pre-stimulus period in this analysis because previous studies have suggested that spontaneous EEG power during this period predicts subsequent confidence in perceptual decisions (Samaha et al., 2017; Wöstmann et al., 2019).

In Figure 5, the direction and strength of relationships between EEG power and confidence, controlling for difficulty level and 1^st^-order accuracy, are represented as t-values averaged across all electrodes at each time and frequency point. These t-values represent group-level tests of whether the regression coefficients (EEG power predicting confidence) from the individual single-trial analyses showed a systematic linear relationship across participants. For the *perceptual* task on day 1 (Figure 5A), we found a single negative cluster to predict confidence ratings, starting ∼0.220s after stimulus onset until 1s and spanning 6-38Hz frequency range (cluster statistic = −32,772, *p* = .0025). Similarly, one negative post-stimulus cluster predicted confidence in the *knowledge* task on day 1 (Figure 5B), beginning at ∼340ms until 1s and spanning 4-32Hz (cluster statistic = −50,344, *p* = .0005). The negative relationships indicate that participants’ confidence ratings were higher on trials with greater alpha and beta desynchronisation (i.e., lower alpha/beta power) following stimulus onset for both tasks. As we observed this effect after controlling for difficulty level and accuracy in the multiple regression models, it indicates that post-stimulus alpha and beta power encode confidence over and above external stimulus information (Griffiths et al., 2019) and objective accuracy. Both the *perceptual* and *knowledge* task effects were globally distributed across the scalp (all 64 electrodes were included in significant clusters), as shown in the topographical maps of mean t-values (Figure 5A, 5B: right inset). Interestingly, these results were only replicated on day 2 for the knowledge but not the perceptual task (Figure 5C-D). In the knowledge task, we found a significant negative cluster beginning at ∼-200ms until 1s and spanning 7-38Hz with all 64 electrodes significant (cluster statistic = −55,987, *p* = .0005), in line with the day 1 results. However, we did not find any significant clusters predicting confidence in the perceptual task on day 2 (Figure 5C: all negative cluster *p*’s => .359). In contrast to previous literature (Samaha et al., 2017; Wöstmann et al., 2019), we found no significant pre-stimulus clusters predicting confidence.

**Figure 5:**
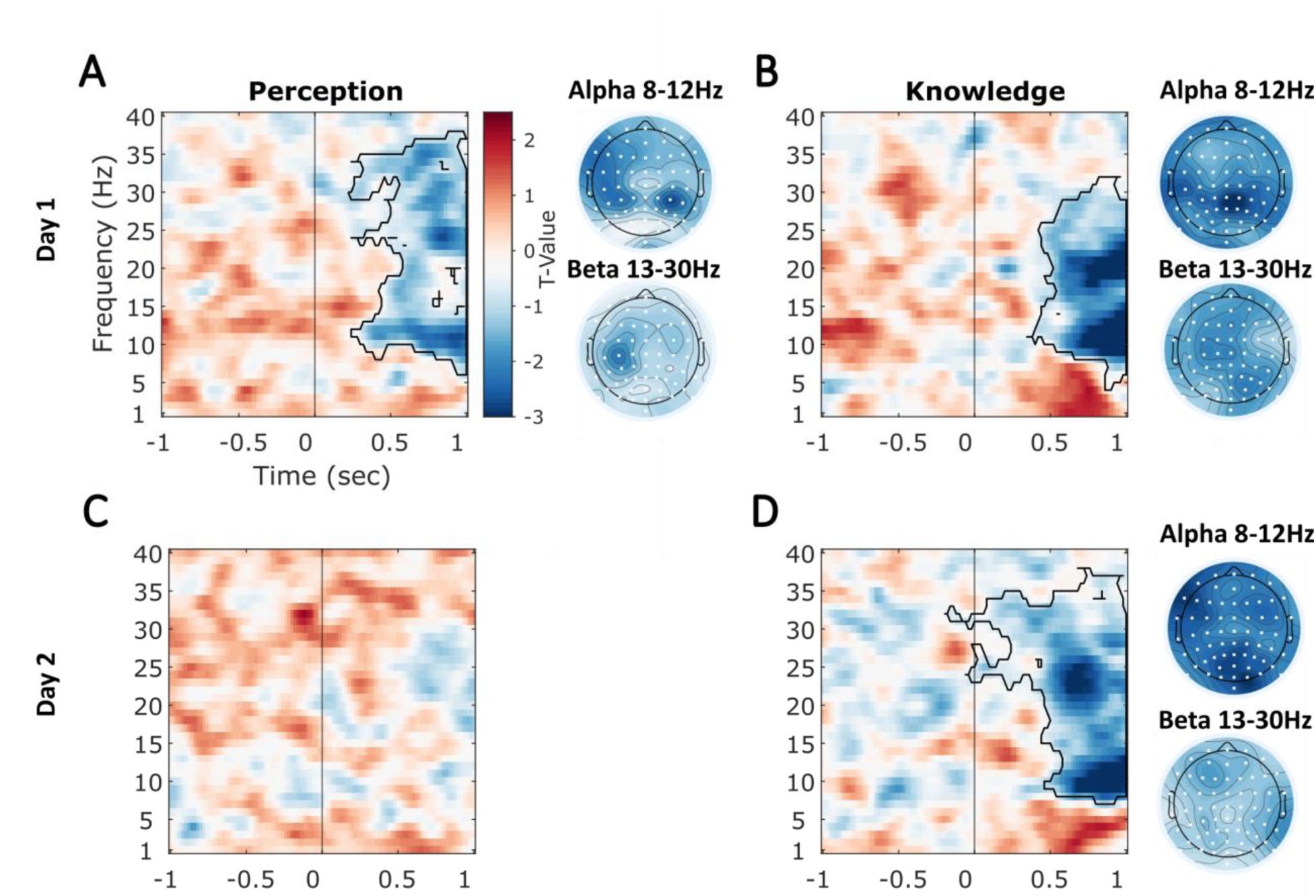
Single-trial alpha/beta desynchronisation reflects subjective confidence ratings, independently of accuracy and evidence discriminability. The time-frequency plots show the mean t-values across all electrodes at each time-frequency point for the **A** Perception task (day 1), **B** Knowledge task (day 1), **C** Perception task (day 2), and **D** Knowledge task (day 2). We obtained the T-values by comparing the single-trial regression coefficients (representing the unique relationship between confidence and spectral power) against zero with cluster-based permutation testing. The significant clusters are outlined in black contour (dependent-samples t-test, 2-tailed, cluster *p* < .025). The black vertical line at 0s denotes stimulus onset. The right inset of each panel plots topographies of the average t-values for the significant clusters across alpha (8-12Hz) and beta (13-30Hz) frequencies separately. Electrodes included in each cluster are highlighted in white.

#### Time-resolved decoding of confidence from ERPs

To further investigate the domain-generality and test-retest reliability of the observed EEG-confidence relationships, we performed additional analyses involving multivariate decoding of confidence from single-trial EEG activity. We reasoned that if a classifier trained to dissociate high versus low confidence trials using single-trial activity on one task could successfully decode confidence on the other (untrained) task, then the neural signature must be engaged during both perceptual and knowledge-based confidence judgements (i.e., it must be domain-general). In contrast, if a signature is domain-specific then the classifier will not be able to predict confidence ratings on the non-trained task. Similarly, if a classifier trained on day 1 could successfully decode confidence on (untrained) day 2, then this would indicate high test-retest reliability of the EEG-confidence signature. Additionally, predictions based on cross-validation analyses constitute stronger evidence than inferential models, as cross-validation directly quantifies the ability to generalise predictions to new data (Bzdok et al., 2020).

First, to establish whether confidence could be decoded from the ERP data we performed within-task decoding of confidence on each testing day separately. We trained and tested LDA classifiers on each point in time (0:1s, stimulus-locked) to investigate how the discrimination pattern generalised over time with a 5-fold cross-validation. In both tasks and on both days, we found significant decoding of confidence. Figure 6 shows significant within-task, within-day clusters beginning at ∼500ms (Fig 6A: day 1, *p* < .0001) and ∼400ms (Fig 6B: day 2, *p* = .001) for the perceptual task. We also trained a classifier on ERP data from day 1 and tested it on data from day 2 (separately for each task) to examine test-retest reliability of the neural markers of confidence. We found significant between-day decoding for the perceptual task beginning at approximately 430ms (Fig 6C: *p* = .0005). In the knowledge task, there were significant within-task, within-day clusters beginning at ∼340ms (Fig 6D: day 1, *p* = .0005) and ∼450ms (Fig 6E: day 2, *p* < .0001). In line with the perceptual task, we found significant between-day decoding for the knowledge task at approximately 430ms (Fig 6F: *p* = .001). For all classifiers, the best decoding was centred around the diagonal – where the training and testing data came from the same time points. However, there was also statistically significant off-diagonal decoding suggesting a degree of temporal generalisation of the classifier discrimination performance. Overall, confidence could be reliably decoded for both tasks over separate days from the late-ERP, which represents a temporally stable neural marker of subjective self-evaluation.

**Figure 6:**
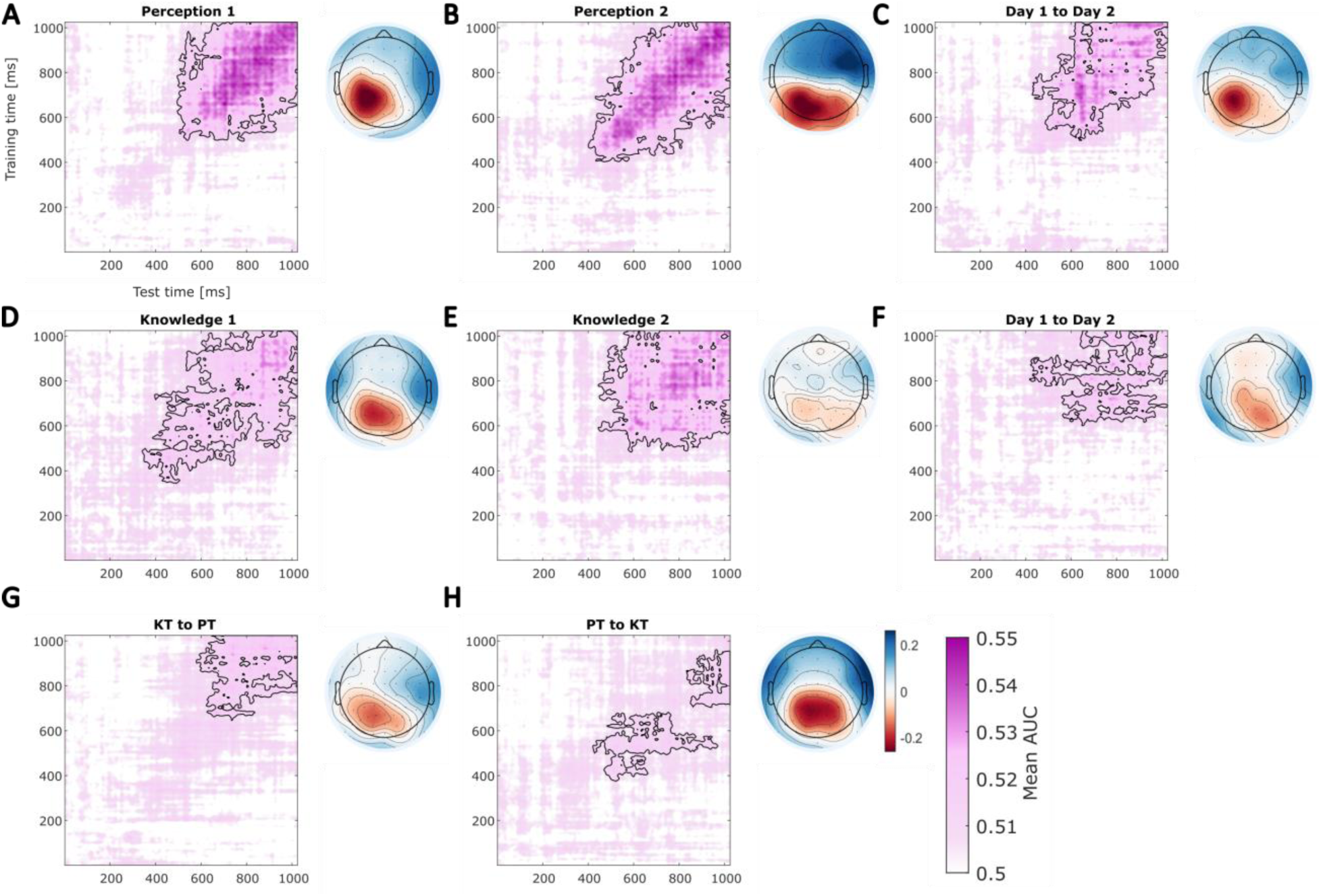
Time-resolved decoding of decision confidence from single-trial stimulus-locked ERPs. Linear Discriminant Analysis (LDA) classifiers were trained and tested at all post-stimulus (0-1s) time points. Mean AUC values across participants are shown. The topographies show group averaged correlations between the classifier decision values and the ERP voltages at each participant’s peak AUC time-point. Note that negative decision values correspond to high confidence trials so that overall negative correlations plotted represent a positive relationship between confidence and ERP amplitude. **A-B** Within-task (**A** day 1, **B** day 2) decoding of perceptual confidence. **C** Cross-day decoding of confidence in perception, where the classifier was trained on data from day 1 and tested on data from day 2. **D-E** Within-task (**D** day 1, **E** day 2) decoding of knowledge confidence. **F** Cross-day decoding in knowledge task. **G-H** Cross-task decoding using data combined across both days. Significant clusters (one-tailed t-test, *p* < .05) are highlighted in black.

To further test the domain-generality of the ERP-confidence relationships, we performed time-resolved cross-task decoding, whereby a classifier was first trained on one task (data combined across both days) and then tested on the data from the other task (and vice versa). We found significant cross-task decoding in both decoding directions (knowledge to perception, and perception to knowledge). For the classifier trained on the knowledge task, we found a significant cluster beginning at ∼580ms (*p* = .0015) (Figure 6G). There were two significant clusters, spanning ∼430-970ms (*p* = .01) and ∼830-1000ms (*p* = .032), respectively, in the performance of the classifier trained on the perceptual task (Figure 6H). These results therefore support the late-ERP as a reliable domain-general predictor of confidence ratings.

#### Time-resolved decoding of confidence from spectral power

Next, we recreated the above analyses using time-frequency spectral power. To improve the signal-to-noise ratio while also being able to examine the temporal generalisability of any EEG power-confidence relationships we performed the decoding using the canonical frequency bands. We focused on alpha (8-12Hz) and beta (13-30Hz) bands based on previous literature showing their link to subjective task performance like perceptual awareness (Benwell et al., 2017, 2022) and confidence (for response-locked activity, Faivre et al., 2018). To test whether confidence can be decoded from alpha + beta activity (combined), we trained and tested LDA classifiers on each post-stimulus time point (0:1s) using a 5-fold cross-validation as described above. We found significant cluster-level decoding in both task and on both days (Figure 7A-B and 7D-E). In the perceptual task, we found significant within-task, within day decoding with a cluster beginning at ∼360ms (*p* < .0001) on day 1 (Figure 7A), and a cluster beginning at ∼600ms (*p* = .0025) on day 2 (Figure 7B). To test the test-retest reliability of these neural signatures we trained a classifier on day 1 data and tested it on day 2 data (separately for each task). Figure 7C shows we did not find any significant clusters in the perceptual task (all *p*’s > .287) decoding confidence across-days. In the knowledge task, we also found significant within-task within day decoding (Figure 7D-E). On day 1 there was a large significant cluster starting at ∼440ms (p < .0001) and a smaller one from ∼60 to ∼400ms (*p* = .042, Figure 7D). On day 2 of knowledge task, we found a cluster spanning ∼440 to ∼980ms, *p* < .0001 (Figure 7E). In contrast to perception, we found significant cross-day decoding in the knowledge task (in line with the results of single-trial regression analyses), with a cluster starting at ∼540ms (*p* = .0005, Figure 7F). When we used power from individual alpha and beta frequency bands to train the classifiers, we obtained similar results, with significant decoding found in both tasks and on both days (apart from alpha power decoding in perceptual task on day 2 which did not reach significance) (see Supplementary Figures S3 & S4). When we included each frequency band in the between-day classification models separately, we found no significant cross-day decoding for 8-12Hz power, while for 13-30Hz power we found significant, though, weaker, decoding in the knowledge task. Hence, this indicates alpha + beta (8-30Hz) spectral power is only a reliable predictor of semantic knowledge confidence.

**Figure 7:**
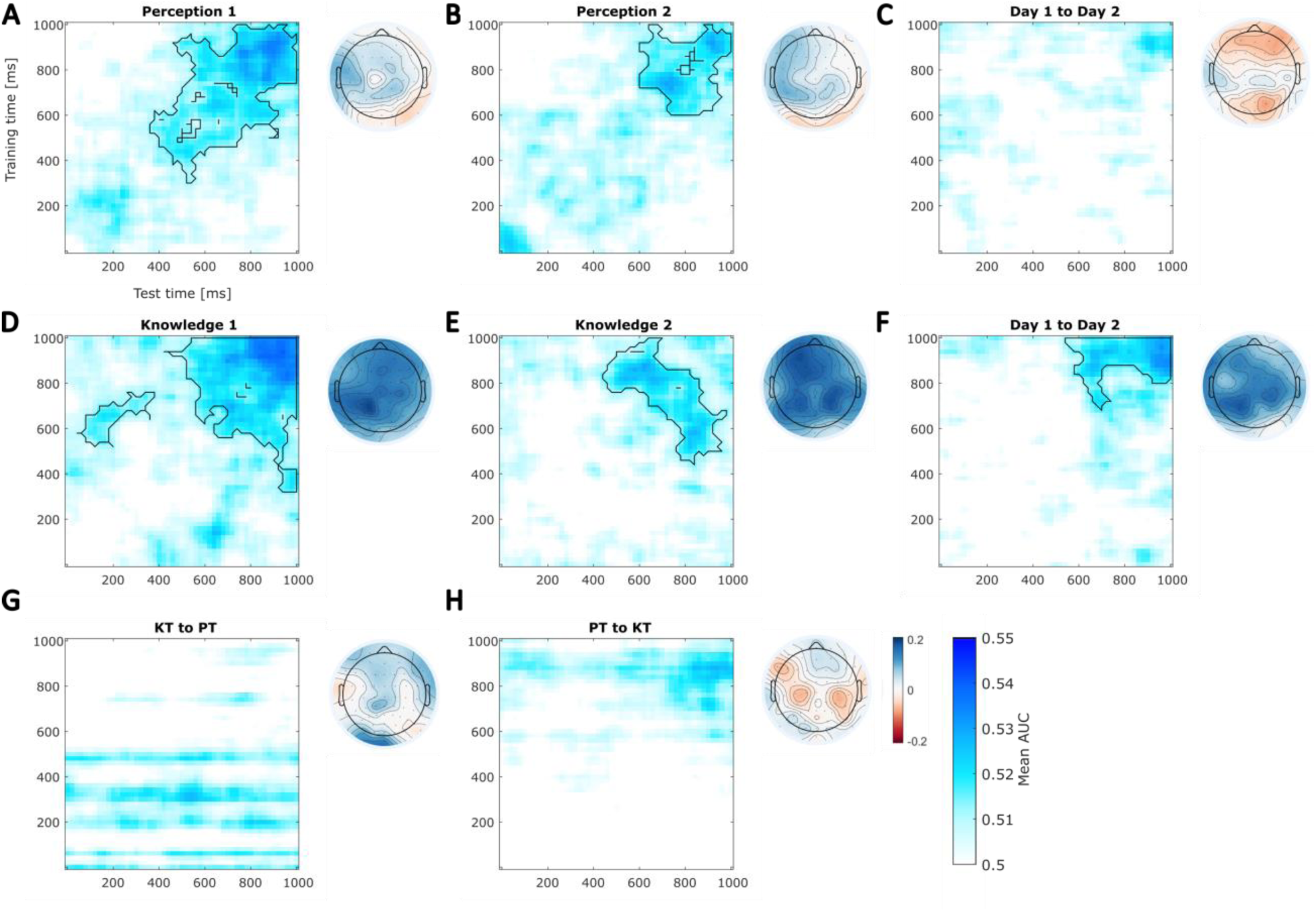
Time-resolved decoding of decision confidence from single-trial α+β (8-30Hz) spectral power (stimulus-locked, 0-1s). The classifiers were trained and tested on all post-stimulus (0-1s) time points. Mean AUC values across participants are shown. The topographies show group averaged correlations between the classifier decision values and the spectral power at each participant’s peak AUC time-point. Note that negative decision values correspond to high confidence trials so that overall positive correlations plotted represent a negative relationship between confidence and 8-30Hz power. **A-B** Within-task (A day 1, B day 2) decoding of perceptual confidence. **C** Cross-day decoding of confidence in perception, where the classifier was trained on data from day 1 and tested on data from day 2. **D-E** Within-task (D day 1, E day 2) decoding of knowledge confidence. **F** Cross-day decoding in knowledge task. **G-H** Cross-task decoding using data combined across both days. Significant clusters (one-tailed t-test, *p* < .05) are highlighted in black.

Finally, we tested whether a classifier trained on one task can predict confidence ratings in the other task to establish the extent of domain-generality. As Figure 7G-H shows, we found no significant cross-task decoding in either decoding direction for 8-30Hz power (knowledge to perception (Fig 7G), nor perception to knowledge (Fig 7H), all *p*’s > .05). Similarly, we found no consistent significant decoding when we tested 8-12Hz and 13-30Hz power separately (Supplementary Figures S3-4).

Overall, these results suggest spectral power in the alpha + beta band (8-30Hz) only reliably predicts confidence in a domain-specific manner, relating more closely to confidence ratings in the knowledge task.

## 4. Discussion

To understand the neurocomputational mechanisms underlying metacognition, it is crucial to establish whether its behavioural and neurophysiological signatures are reliable over time and generalisable across cognitive domains. Here, we examined the test-retest reliability and domain-generality of both behavioural and EEG measures of metacognition. Overall confidence calibration (i.e., metacognitive bias) was highly domain-general across two cognitive domains (perception and semantic memory) and showed strong test-retest reliability across two experimental sessions. In contrast, metacognitive efficiency (indexed by meta-d’/d’) showed low domain-generality and poor test-retest reliability. These results suggest that both domain-general and domain-specific resources contribute to overall metacognitive performance, with a dissociation between metacognitive bias and efficiency, wherein the former may be underpinned by more global metacognitive mechanisms. Trial-by-trial fluctuations of confidence within-participants were predicted by both stimulus-locked ERP and ERSP activity during decision formation, though the ERP signature was the most reliable predictor across domains and sessions. The results 1) reveal the test-retest reliability of the most widely adopted measures in the field, 2) contribute importantly to the debate about domain-generality of metacognition, and 3) show that confidence is encoded in two distinct stimulus-locked EEG signatures during decision formation.

### Test-retest reliability and domain-generality of metacognitive measures

High test-retest reliability of model-based measures is essential to build a robust science of metacognition and for translating findings into relevant fields such as computational psychiatry (Brown et al., 2020). To our knowledge, this is the first study to report the test-retest reliability of one of the most widely adopted measures of metacognitive efficiency (meta-d’/d’, often called the M-ratio (Fleming & Lau, 2014; Maniscalco & Lau, 2012)). For both the perception and knowledge tasks, the M-ratio showed poor test-retest reliability. This extends previous research showing poor split-half reliability of M-ratio within a single testing session (Guggenmos, 2021). The M-ratio was also not strongly correlated between tasks, in line with studies that found no (or weak) domain-generality of efficiency measures (Ais et al., 2016; Baird et al., 2013; Benwell et al., 2022b; Fitzgerald et al., 2017). Metacognitive efficiency was significantly higher for knowledge than perception, and this effect replicated across days despite the low test-retest reliability of the individual participant measures, while type-1 sensitivity (d’) was higher for perception. It remains to be seen whether the lack of reliability across time and domains represents inherent variability of metacognitive efficiency itself (Bang et al., 2019; Shekhar & Rahnev, 2021a) or noisiness of the M-ratio measure. Alternative efficiency measures have recently been proposed (Desender et al., 2022; Guggenmos, 2022; Miyoshi & Nishida, 2022; Shekhar & Rahnev, 2021a), and their test-retest reliability and domain-generality should be investigated in future studies.

In contrast to metacognitive efficiency, confidence bias (type-2 c’) was highly reliable between testing sessions in both cognitive domains and showed strong domain-generality. This is in line with previous studies that found strong between-task correlations of metacognitive bias (Ais et al., 2016; Benwell et al., 2022b; Mazancieux et al., 2020), suggesting overall confidence level is a domain-general, trait-like characteristic that is less influenced by task-specific representations and evidence type. This highlights that maladaptive confidence calibration may have a more pervasive influence on everyday life, in line with its association with symptoms of psychopathology (Benwell et al., 2022b; Rouault, Seow, et al., 2018) and longitudinal evidence of confidence bias improvement with mental health treatment (Fox et al., 2023). Hence, identifying reliable, domain-general EEG predictors of confidence may represent a first step towards understanding neural mechanisms relevant for mental health.

### CPP/P3 is a reliable and domain general predictor of confidence

Across both tasks and both sessions, within-participant fluctuations in the level of confidence from trial-to trial were correlated with a late ERP component, likely corresponding to the Central Parietal Positivity (CPP)/P300 based on its latency and topography (Kelly & O’Connell, 2013; O’Connell et al., 2012; Tagliabue et al., 2019). Higher amplitudes of this component during decision formation preceded higher confidence ratings. This is consistent with previous studies which found that the CPP scales with both implicit (e.g., statistical) and explicit confidence level (Feuerriegel et al., 2022; Fitzgerald et al., 2022; Gherman & Philiastides, 2015, 2018; Grogan et al., 2023; Herding et al., 2019; Rausch et al., 2020; Zakrzewski et al., 2019). However, previous studies have focused predominantly on perceptual confidence. Here we show that the CPP/P300 also tracks explicit post-decisional reports of confidence, over and above trial difficulty and accuracy, even when internal (rather than sensory) evidence is evaluated during semantic knowledge decisions. Moreover, we were able to train a classifier that successfully discriminated between high/low confidence trials across testing days and cognitive domains, hence providing strong evidence for test-retest reliability and domain-generality of this neural signature. The test-retest reliability of the effect is particularly important given increasing concerns regarding the reliability of scientific findings over the last decade, with numerous replication failures, particularly within neuroscience (Botvinik-Nezer et al., 2019; Haines et al., 2023; Pavlov et al., 2021; Poldrack et al., 2017).

The stimulus-locked CPP, which closely matches our confidence discriminating ERP component, is thought to reflect a supramodal decision signal that tracks accumulating evidence (Kelly & O’Connell, 2013; O’Connell et al., 2012). Hence, the relationship with confidence suggests that 1^st^ and 2^nd^-order decision related signals may unfold together during decision formation, in line with these processes being coupled in the early stages of decision formation (Gherman & Philiastides, 2015; Kiani & Shadlen, 2009; van den Berg et al., 2016). Interestingly, Gherman and Philistines (2018) identified a confidence discriminating component, similar in time and topographical distribution to ours, which simultaneous fMRI revealed originated in ventromedial Prefrontal Cortex (vmPFC). The vmPFC has also been shown to encode confidence in other functional neuroimaging studies (De Martino et al., 2013; Lebreton et al., 2015; Morales et al., 2018). Hence, we speculate that the confidence ERP effect we observed may reflect a domain-general PFC metacognitive signal.

### Alpha/beta de-synchronisation also predicts confidence

In addition to the ERP-confidence relationship, greater post-stimulus alpha/beta desynchronisation also predicted higher confidence ratings, over and above accuracy and trial difficulty, in both the perceptual and knowledge tasks. Post-stimulus alpha/beta desynchronisation has previously been shown to correlate with perceptual awareness (Benwell et al., 2017, 2022a) and perceptual confidence (Faivre et al., 2018). Here we show that alpha/beta desynchronisation is associated with explicit confidence ratings during decision formation across both perceptual and semantic knowledge decisions. However, the effect was only reliable over time in the knowledge task (and not perception), suggesting it is more domain-specific than the ERP effect. Alpha desynchronisation is proposed to reflect increases in cortical excitability modulated by global noradrenergic arousal according to attentional demands (Dahl et al., 2022; Kosciessa et al., 2021). Importantly, higher confidence ratings have also been related to physiological measures of arousal like increased heart rate (Allen et al., 2016). Hence, it is possible that the fluctuations in post-stimulus power we observed reflect internal states, such as attention and arousal, that concurrently influence confidence. This could be tested in the future by combining EEG recordings with other physiological measures. Additionally, the global alpha/beta desynchronisation effects we observed could result from oscillatory communication in widespread neural networks (Buzsáki & Draguhn, 2004). The lack of successful cross-task decoding may suggest that although alpha/beta mechanisms are linked to confidence in both tasks, the networks involved, and hence the topographical patterns captured in our decoding analysis, are relatively domain-specific.

Importantly, both late ERPs and alpha/beta desynchronisation correlated with confidence independently of evidence strength and 1^st^-order accuracy. Dissociation between subjective confidence and objective performance is in line with 2^nd^-order confidence generation models (Fleming & Daw, 2016; Pleskac & Busemeyer, 2010) which propose that, unlike accuracy, confidence may rely on additional computations (Murphy et al., 2015) and be more susceptible to factors beyond evidence strength, like attention (Denison et al., 2018). Our results suggest that prior to the 1^st^-order response these computations are linked to both decision-related ERPs as well as alpha/beta power, whereby the former may represent a metacognitive readout of accumulating decision evidence (Gherman & Philiastides, 2018) while the latter may be more closely linked to additional influences such as arousal and/or attention (Dahl et al., 2022; Kosciessa et al., 2021).

It is important to note that other EEG signatures have been linked to confidence such as the response-locked error-related negativity and Error Positivity (Boldt & Yeung, 2015; Desender et al., 2019, though see Feuerriegel et al., 2022; Grogan et al., 2023; Rausch et al., 2020). Here, we focused on stimulus-locked activity and pre-response neural signatures of confidence by delaying the response period by 1s from stimulus onset. Hence, we were unable to conduct meaningful reaction time or response-locked analyses. Future studies can investigate the reliability and domain-generality of these alternative confidence signatures.

In conclusion, confidence calibration (metacognitive bias) is highly reliable across time and cognitive domains. In contrast, metacognitive efficiency (as indexed by meta-d’/d’) is not reliable over time, nor between tasks. This emphasises the need for the development of more reliable model-based measures of metacognition that may need to be tailored according to different types of decision evidence. We show that two distinct neural signatures encode confidence judgements: slow ERPs which are reliable across both time and cognitive domains, and alpha/beta desynchronisation which is stronger and more reliable across time in the semantic knowledge domain (relative to perception). Identifying reliable and domain-general neural predictors of confidence represents a crucial step towards understanding the neural mechanism underlying suboptimal self-evaluation.

## Acknowledgments

C.S.Y.B. is supported by the British Academy/Leverhulme Trust and the United Kingdom Department for Business, Energy and Industrial Strategy [SRG19/191169]. M.K. is supported by the United Kingdom Economic & Social Research Council Scottish Graduate School of Social Science [ES/P000681/1]. The authors thank Dewy Nijhof for help with PsychoPy.

