## Supplementary Figure for "Two distinct stimulus-locked EEG signatures reliably encode domain-general confidence during decision formation"

Abbreviated title: Reliability and domain-generality of metacognition.

Martina Kopčanová, Robin A. A. Ince[2], Christopher S. Y. Benwell[1]

[1] Division of Psychology, School of Humanities, Social Sciences, and Law, University of Dundee, Dundee, DD1 4HN, UK

[2] School of Psychology and Neuroscience, University of Glasgow, Glasgow, G12 8QB, UK

Corresponding authors: Christopher S. Y. Benwell & Martina Kopčanová

Division of Psychology, School of Humanities, Social Sciences, and Law, University of Dundee, Dundee, UK

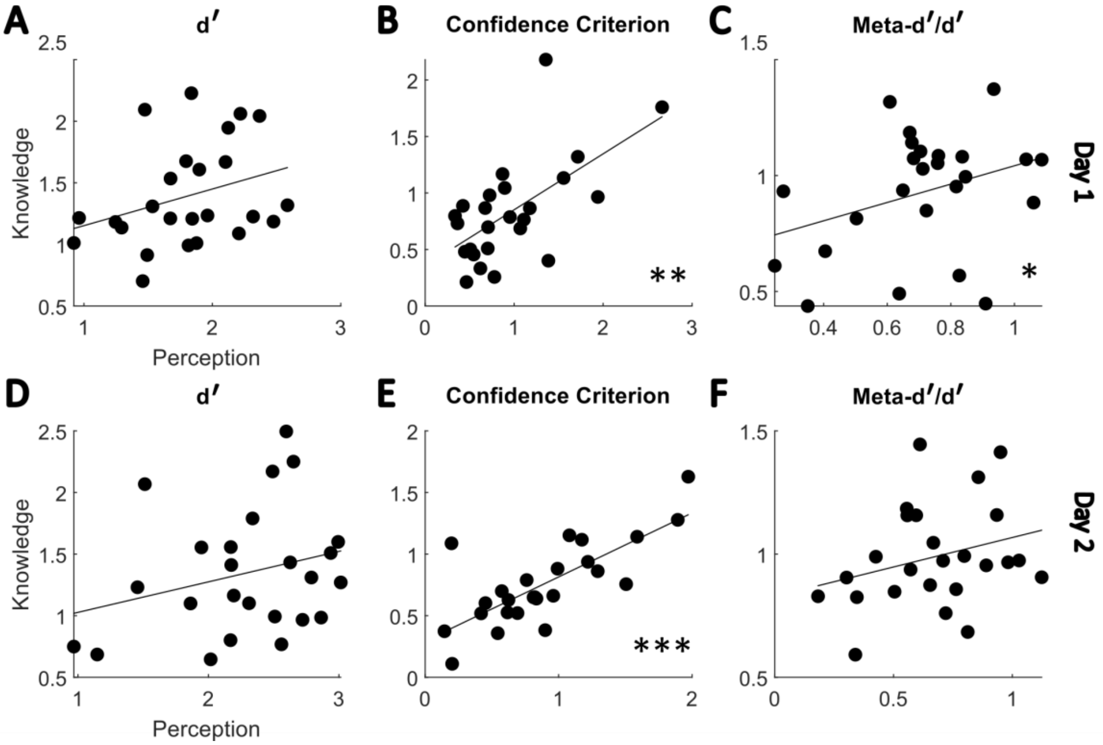


**Figure S1** Between-task correlations for each model-based measure. The scatterplots correspond to the ICC results reported in Table 1 in the main text. **A-C** Day 1 between-task relationships for type-1 sensitivity (d'), confidence bias (type-2 (confidence) criterion), and metacognitive efficiency (M-ratio). **D-F** Day 2. *** $p$ < .0001, ** $p$ < .001, * $p$ < .05

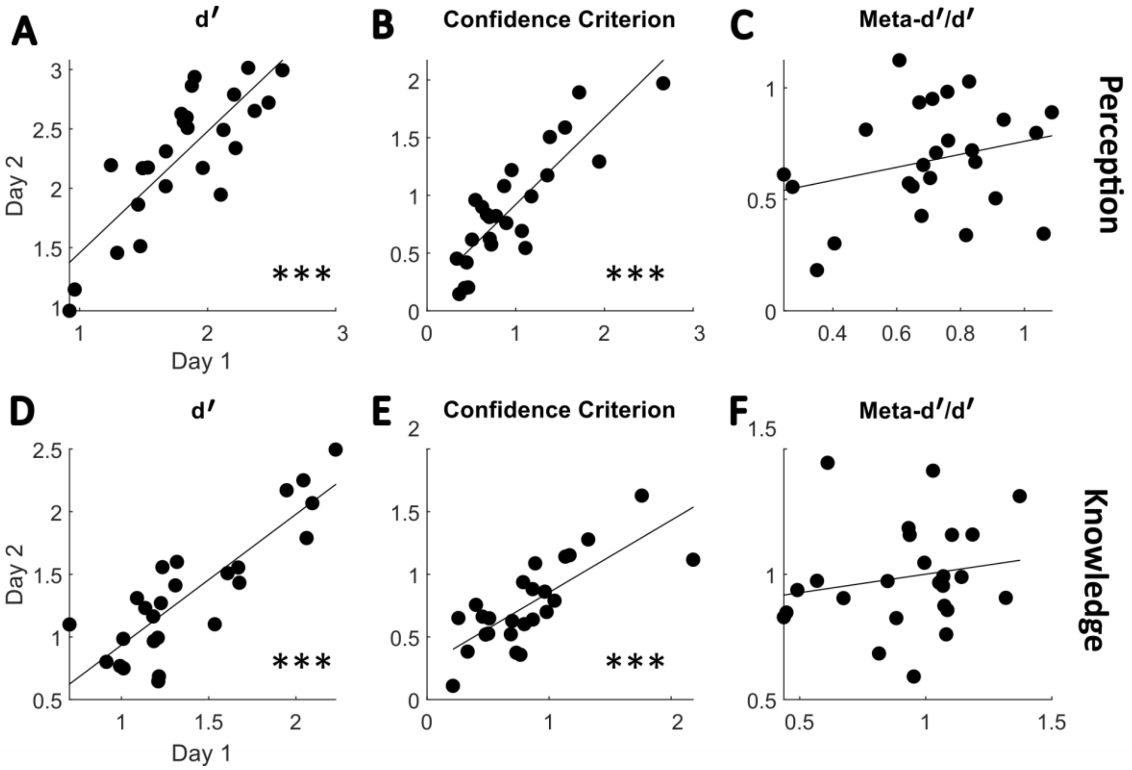


**Figure S2** Between-day correlations for each model-based measure. The scatterplots correspond to the ICC results reported in Table 1 in the main text. **A-C** Perceptual task between-day relationships for type-1 sensitivity (d'), confidence bias (type-2 (confidence) criterion), and metacognitive efficiency (M-ratio). **D-F** Knowledge task. *** $p < .0001$, ** $p < .001$, * $p < .05$

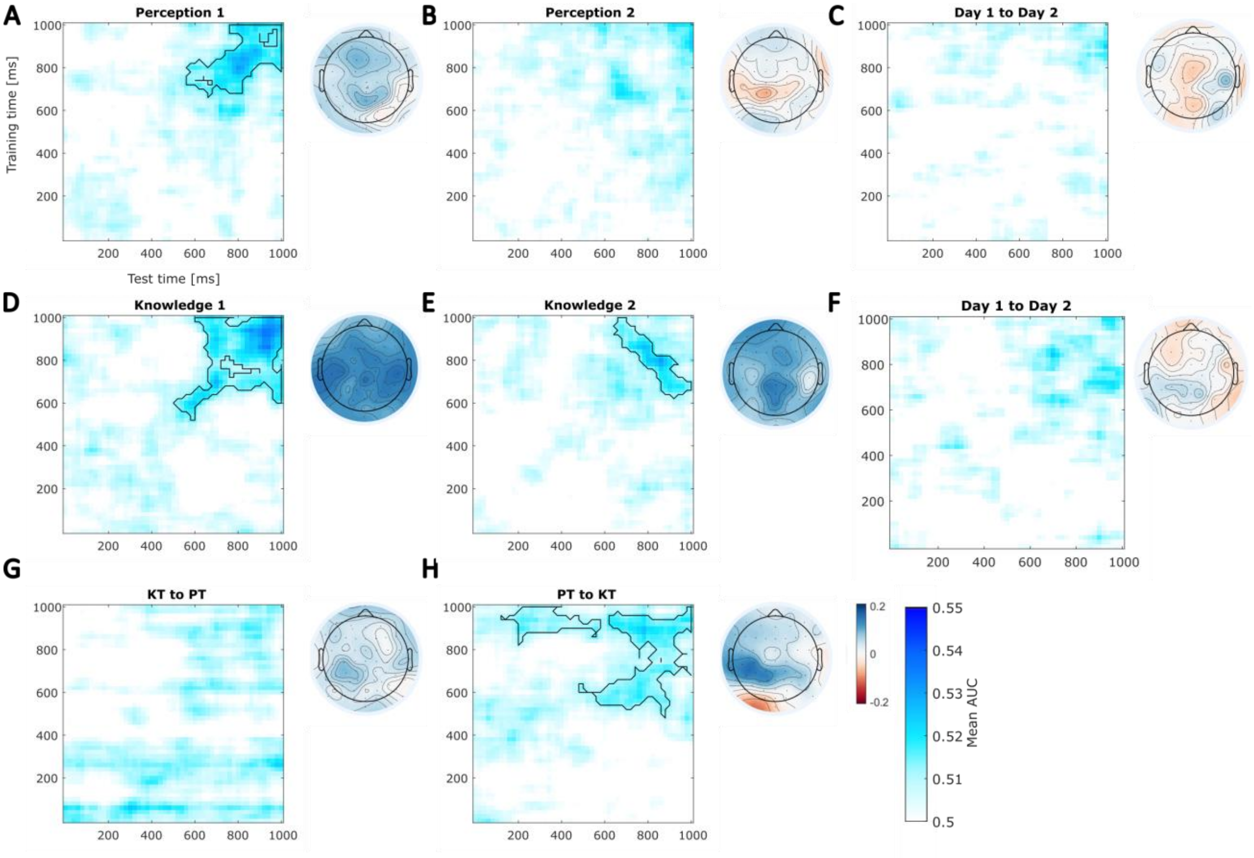


**Figure S3** Time-resolved decoding of decision confidence from single-trial α (8-12Hz) spectral power (stimulus-locked, 0-1s). The classifiers were trained and tested on all post-stimulus (0-1s) time points. Mean AUC values across participants are shown. The topographies show group averaged correlations between the classifier decision values and the spectral power at each participant's peak AUC time-point. Note that negative decision values correspond to high confidence trials so that overall positive correlations plotted represent a negative relationship between confidence and 8-12Hz power. **A-B** Within-task (A day 1, B day 2) decoding of perceptual confidence. **C** Cross-day decoding of confidence in perception, where the classifier was trained on data from day 1 and tested on data from day 2. **D-E** Within-task (D day 1, E day 2) decoding of knowledge confidence. **F** Cross-day decoding in knowledge task. **G-H** Cross-task decoding using data combined across both days. Significant clusters (one-tailed t-test, $p < .05$) are highlighted in black.

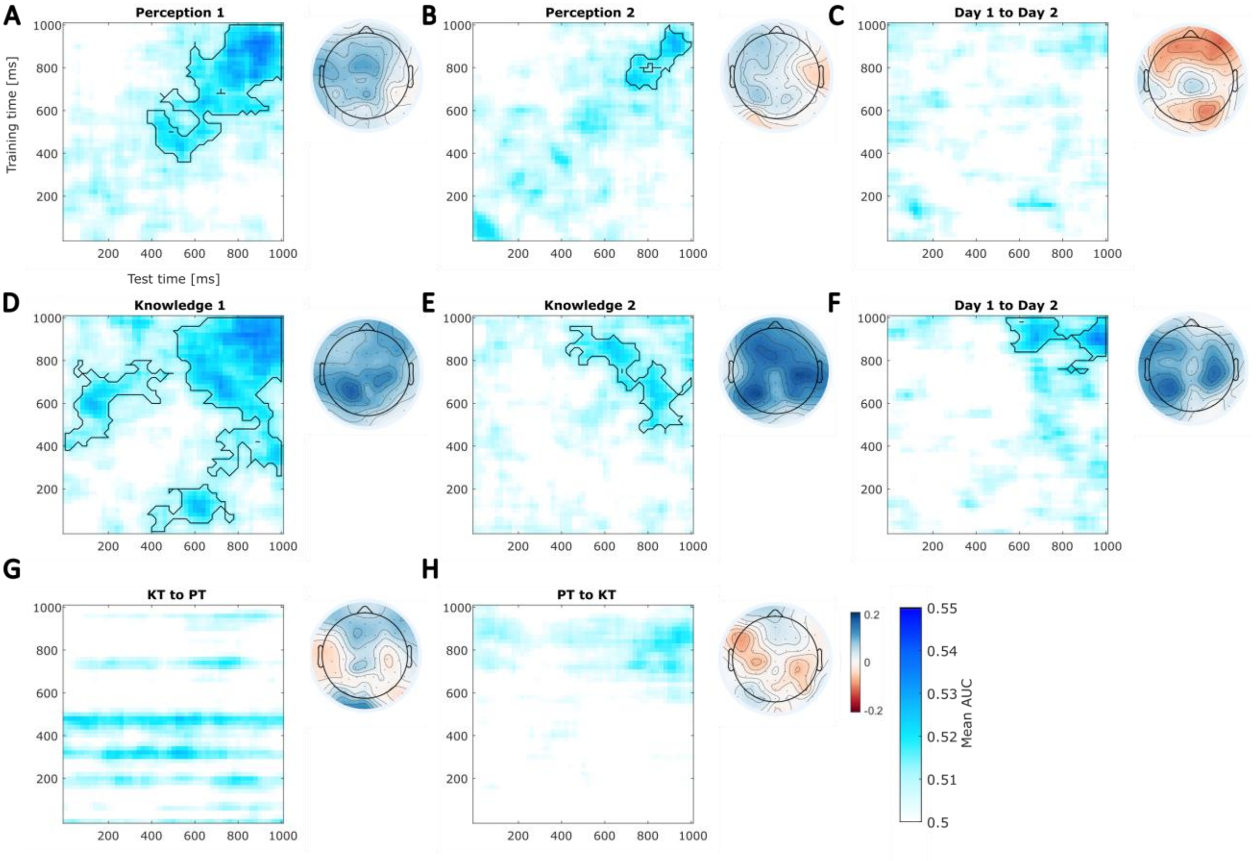


**Figure S4** Time-resolved decoding of decision confidence from single-trial β (13-30Hz) spectral power (stimulus-locked, 0-1s). The classifiers were trained and tested on all post-stimulus (0-1s) time points. Mean AUC values across participants are shown. The topographies show group averaged correlations between the classifier decision values and the spectral power at each participant's peak AUC time-point. Note that negative decision values correspond to high confidence trials so that overall positive correlations plotted represent a negative relationship between confidence and 13-30Hz power. **A-B** Within-task (A day 1, B day 2) decoding of perceptual confidence. **C** Cross-day decoding of confidence in perception, where the classifier was trained on data from day 1 and tested on data from day 2. **D-E** Within-task (D day 1, E day 2) decoding of knowledge confidence. **F** Cross-day decoding in knowledge task. **G-H** Cross-task decoding using data combined across both days. Significant clusters (one-tailed t-test, $p < .05$) are highlighted in black.
